# Deciphering the role of the lncRNA TRIBAL in hepatocyte models

**DOI:** 10.1101/2025.04.02.646870

**Authors:** Sébastien Soubeyrand, Paulina Lau, Ruth McPherson

## Abstract

We recently reported that the long non-coding RNA *TRIBAL/TRIB1AL* was required to sustain key hepatocyte functions. Here, we identify HepaRG cells as a model for studying *TRIBAL* and provide additional validation and functional insights. In contrast to HepG2 and HuH-7 cells, differentiated HepaRG cells showed similarities to primary hepatocytes in response to *TRIBAL* suppression. *TRIBAL* suppression was associated with reduced HNF4A and MLXIPL abundance in hepatocytes and HepaRG cells. *TRIBAL* targeting using a panel of targeting ASOs confirmed specificity. Comparing *TRIBAL*-suppressed hepatocyte and HepaRG transcriptomics identified extensive functional overlap. Biological ontologies associated with key hepatic metabolic functions were predicted to be inhibited in both models. Comparative analyses with TRIB1-suppressed HepaRG cells, a major metabolic regulator vicinal to *TRIBAL*, also revealed extensive functional congruence with *TRIBAL*. Interestingly, *TRIBAL* transduction failed to restore function in *TRIBAL*-suppressed cells, which may be linked to structural differences, as supported by contrasting RNAseR sensitivities between the endogenous and transduced forms. In summary, these findings support the use of HepaRG cells as an experimental model to study *TRIBAL* and underscore its importance in regulating key hepatocyte genes essential for metabolic function.

## Introduction

The 8q13 chromosomal region is associated with plasma lipid levels, hepatic steatosis, and the risk of coronary artery disease (CAD). Initially attributed to its gene neighbor *TRIB1*, a major regulator of lipid homeostasis, we recently demonstrated that the genetic correlation was driven proximally by the gene encoding the long non-coding RNA (lncRNA) *TRIBAL,* also known as *TRIB1AL*[1]. Initially dismissed as biological noise, it is now recognized that lncRNAs can regulate gene expression at both the transcriptional and post-transcriptional levels. Using embryonic stem cells and shRNA approaches, Guttman et al. showed nearly 10 years ago that most lncRNAs were likely functional.[2] Since then, the list of lncRNA genes, which probably outnumber protein-coding genes, has increased considerably, although very few are sufficiently characterized [3,4].

We recently demonstrated that suppression of *TRIBAL* in primary hepatocytes potently reduced the expression of several major lipid regulators [1]. However, the underlying mechanisms were not explored due to technical limitations, primarily the human-specific expression of *TRIBAL* and the lack of a suitable model system. Indeed, the limited supply, high cost, and inter-donor variations of primary hepatocytes pose significant challenges, necessitating the need for alternative models. We previously showed that the commonly used immortalized hepatocyte-like cell models, namely HepG2 (hepatoblastoma) and HuH-7 (hepatocarcinoma), were largely unresponsive to *TRIBAL* targeted intervention, specifically *TRIBAL* overexpression (in HepG2) and *TRIBAL* suppression (in HuH-7) [5].

Here, we continue our investigation into *TRIBAL*. First, we identify HepaRG cells as a model system amenable to *TRIBAL* interrogation. HepaRG is a human cell line that can be maintained for several passages and differentiated *in vitro* into a mixture of hepatocyte- and biliary-like cells under appropriate cell culture conditions [6,7]. Although not identical to primary hepatocytes, HepaRG cells more closely resemble hepatocytes as compared to HepG2s [8,9]. We now demonstrate that, similar to primary hepatocytes, *TRIBAL* expression was required to support the expression of major hepatic regulators in HepaRG cells. Specificity was ascertained with several antagomirs. *TRIBAL* suppression reduced HNF4A and MLXIPL protein abundance in primary hepatocytes and HepaRG cells. Transcriptomic datasets from *TRIBAL*-suppressed HepaRG cells were generated with microarrays and compared with the corresponding hepatocyte data, revealing extensive similarities. Comparison of *TRIB1*- and *TRIBAL-*suppressed HepaRG cells also revealed substantial functional convergence. Lastly, we provide evidence that the native *TRIBAL* transcript is resistant to RNaseR digestion, in this regard differing fundamentally from the transduced transcript.

## Methods

### Cell culture and treatments

HepaRG cells were obtained from BIOPREDIC International. Cells were seeded at 29,000 cells per cm^2^ and maintained for 2 weeks in proliferation media, followed by 2 weeks in differentiation media. Initially, proliferation media and differentiation media were obtained from BIOPREDIC (results shown in Figure 2). In the other experiments, homemade growth media, consisting of William’s E medium supplemented with 10% fetal bovine serum, insulin (0.15 U/ml; Humalog, Eli Lilly), Hydrocortisone 21-hemisuccinate (54 µM; Cayman Chemical), and Penicillin-Streptomycin (Gibco), was used, similar to that described previously [6]. Differentiation by inclusion of 1% DMSO for 48 h and of 2% DMSO for an additional 10-14 days. While side-by-side comparisons were not performed, cells grown in the BIOPREDIC media provided greater sensitivity to the *TRIBAL* ASOs (e.g ASO2 in Fig 2 vs Fig 4), which could reflect differences in serum (which are tested by BIOPREDIC to confer optimal differentiation capacity) or proprietary additives leading to better terminal differentiation, a process still poorly understood. Importantly, responses were qualitatively similar (i.e.general suppression of the TOI). Media was changed every 48-72 hours. HuH-7 (Japanese Collection of Research Bioresources Cell Bank) and HepG2 (ATCC) were maintained in DMEM supplemented with 5 mM glucose, 10% fetal bovine serum, and Antibiotic– Antimycotic (Gibco). Cryopreserved primary hepatocytes (HMCPMS), media, and media supplements were obtained from ThermoFisher Scientific. Lots HU8413 (lot 2) and HU8412 (lot 1) were thawed in thawing media and seeded in 12-well plate wells (14-16 wells per vial). The media was changed to a culture medium, which was replaced every 24 hours until the transfection, where incubation was continuous in the presence of the antagomirs. Experiments were initiated 4-6 days post-plating, resulting in 3 distinct replicates for each donor, as described previously[1].

### Viral particle generation

Viral particles were generated in 293FT cells with the empty PLVX plasmid, PLVX*TRIBAL*, psPAX2, and pMD2G. Viral particles harvested during the 16-72 h window post-transfection were concentrated with Lenti-X (Clontech) and stored at - 80°C. Titers were determined on HEK293T cells using puromycin selection (3 µg/ml). Cells were inoculated with three multiplicities of infection (MOI).

### Western blotting

Cells were extracted for 5 min in lysis buffer (50 mM Tris-HCl, 150 mM NaCl, 1% NP40, pH 7.4) supplemented with protease and phosphatase inhibitors (PhoStop and Complete, Roche). Samples were centrifuged (2 min @ 16 kg), and the supernatant (20 µg of protein) was denatured in 1X Laemmli SDS-PAGE buffer (95 °C, 5 min). Samples were then subjected to SDS-PAGE (10% gels) and transferred using liquid transfer (1 h, 90 V) to nitrocellulose membranes. Even loading and transfer were ascertained by Ponceau staining. Blots were then destained and blocked in Intercept buffer (LI-COR) for one hour. Detection was performed using antibodies diluted in PBS/0.1% Tween. Primary antibodies (1:1000 dilution) were incubated for 16 hours, and secondary antibodies (LI-COR; 1:20,000 dilution) for 1 hour. Four 30-second washes in PBS were performed after each antibody incubation. Blots were imaged on an Odyssey Imager (LI-COR). All images were within the instrument’s dynamic range and were only adjusted in contrast and intensity.

### CRISPR and dCAS9 targeting of *TRIBAL*

Pairs of oligonucleotides matching two regions flanking exon one were inserted downstream of the U6 promoter of a trcrRNA expression plasmid (pCRU6)[10]. Cells were transfected with pCRU6 constructs and eSpCas9 a gift from Feng Zhang (Addgene plasmid # 71814) [11]. Transfections were performed in 24-well plate wells using 0.5 μg of DNA with an 8:1:1 (eSpCas9:pCRU6g1:pCRU6g2) DNA mass ratio.

HepG2 and HuH-7 (40-60% confluent) cells were transfected using Extreme Gene HP (3:1, HP: DNA) or Lipofectamine 3000 (3:2:1, Lipofectamine 3000: P3000: DNA). dCAS9 activation was performed using a dual expression plasmid (pAC154-dual-dCas9VP160-sgExpression) encoding inactive CAS9 coupled to VP160 and a U6-driven sgRNA cassette, a gift from Rudolf Jaenisch obtained via Addgene (plasmid # 48240)[12]. Oligonucleotides targeting two regions within the *TRIBAL* promoter were cloned in the U6 cassette. HepG2 and HuH-7 cells were transfected for 72 h with the dCas9VP160 containing sgRNA4, sgRNA5, or trcrRNA control. Oligonucleotide sequences are listed in **Supplemental Material.**

### Antagomir treatment

Antagomirs were designed as gapmers. Sequences are listed in **Supplemental Material.** HepG2 and HuH-7 cells were transfected at 30%-50% confluence in 0.5 mL of media in 24-well plates with 10 nM ASO (final concentration) and 1 µL of RNAiMax. The siRNA transfection mix was prepared in 100 µL of Opti-MEM (Gibco). For HepaRG (24 well-plates, 0.5 ml per well) and primary hepatocytes (12 well-plates, 1 ml per well), transfection was performed using 60 nM ASO (final) and 2.4 µl (HepaRG) or 4.8 µl (hepatocytes) of RNAiMax in 100 µl (HepaRG) or 200 µl (hepatocytes) Optimem. Treatment was continuous for 72 hours.

### Real-Time RNA Quantification PCR (RT-qPCR)

RNA was extracted from culture plates using TriPure Reagent (Roche) and isolated using Direct-Zol RNA mini prep kits (Zymo Research). RNA (0.25 µg in 10 µl) was reverse-transcribed using the Transcriptor First Strand cDNA kit (Roche), employing a 1:1 mixture of oligo dT and random hexamer primers for 1 hour. The resulting cDNA was diluted sixfold in H2O and quantified using a Light Cycler 480 with SYBR Green (Roche) and 0.5 µM primers in 384-well plates. Target of interest values were expressed relative to the corresponding peptidyl-prolyl isomerase A (PPIA) values using the ΔΔCt method. PPIA is routinely used in our laboratory as a robust housekeeping gene in hepatocyte models based on its insensitivity to *TRIBAL* (and TRIB1) ASO treatments in qRT-PCR and transcription array expression datasets (e.g., GSE284599, GSE248931, GSE61473). Oligonucleotides are detailed in the “**Supplemental material”** section.

### RNAse R digestion

Total RNA was purified using Tri-Reagent (Roche) and Direct-Zol RNA miniprep kits (Zymo Research). RNA (500 ng) was then incubated with 0.5 µl of RNase R (Ambion) for 2 hours at 37°C in a total volume of 25 µl. Samples were then reverse-transcribed as described above and analyzed by qPCR. Relative sensitivity was calculated using the ΔΔCt method by comparing the abundance of mock-incubated RNA to the matching RNase R-treated value.

### Transcriptome clustering analyses

Three transcription array datasets consisting of antagomir-treated Hepatocytes and HepaRG cells were used: GSE61473 (Hepatocytes, TRIB1ASO1, 48 h), GSE248931 (Hepatocytes, *TRIBAL*ASO1 and ASO2, 72 h), and GSE284599 (HepaRG, *TRIBAL* ASO2 and *TRIB1* ASO2, 72 h). Data are available at the Gene Expression Omnibus depository. The entire list of mappable Entrez Gene IDs was used for GSEA, using the Gene Ontologies (Biological noRedundant), through WebGestalt. Rather than relying on an arbitrary significance threshold (as in overrepresentation analyses), GSEA employs a ranking approach that aggregates incremental changes in transcripts within categories to better capture enrichment patterns within a dataset, compared to overrepresentation analysis. Output was limited to a maximum of 200 terms, and highly significant (p < 0.01) terms were retained. Comparative analyses were conservatively performed on FDR-significant terms to minimize false positives. Thus, the actual overlap is probably underestimated. The clustering of enriched GO terms based on semantic similarity was performed using Revigo, using the default parameters [13]. Visualization was performed via Cytoscape[14].

## Results

### The *TRIBAL* locus, but not its transcript, is required for optimal growth of HepG2 and HuH-7 cells

*TRIBAL* suppression in primary hepatocytes resulted in pervasive transcriptome-wide changes in primary hepatocytes[1]. Earlier efforts using an ASO (ASO1) to suppress *TRIBAL* in HuH-7 and *TRIBAL* overexpression in HepG2 cells failed to uncover consistent transcriptome-wide changes [5]. Examination of a subset of transcripts of interest (TOIs), selected based their response to *TRIBAL* suppression and functional importance in hepatocytes, revealed little to no consistent effects in response to suppression using two distinct *TRIBAL*-targeting ASOs (**Fig. 1**). Similarly, increased *TRIBAL* expression via CRISPR activation did not affect the TOIs (**Fig. S1**). Thus, these model systems were unsuitable for shedding light on the role of TRIBAL, despite the locus’s importance in these cell lines, as demonstrated by CRISPR deletions, which indicated that the locus was under selective pressure (**Fig. S2**).

### The HepaRG model partially recapitulates the dependence of primary hepatocytes on *TRIBAL*

Looking for a suitable cell model alternative, HepaRG cells were examined next. HepaRG cells acquire hepatocyte-like gene expression and morphology upon differentiation and resemble hepatocytes more than HepG2 cells [8,15,16]. Unlike HepG2 and HuH-7 cells, *TRIBAL* suppression in differentiated HepaRG cells reduced the expression of targets of interest previously shown to be impacted in primary hepatocytes **(Fig. 2)** [1]. As observed previously in primary hepatocytes, suppression was particularly evident with ASO2, although a concordant but significantly weaker pattern was observed with ASO1. Moreover, *HNF4A* expression was also reduced, consistent with our previous finding that HNF4 function is impaired in *TRIBAL*-suppressed hepatocytes. Immunoblotting confirmed the reduction in HNF4A and MLXIPL protein abundance in both primary hepatocytes and HepaRG, although the decrease in HepaRG was observed only with ASO2 (**Fig. 3**).

### Identification of additional *TRIBAL*-suppressing antagomirs

Next, we examined whether the stronger suppressive ability of ASO2 (compared to ASO1), also observed in hepatocytes, resulted from possible off-targeting, by identifying additional *TRIBAL*-targeting ASOs. After an exploratory screen of 5 additional ASOs in HepaRG, 4 showed evidence of suppression (2 vs. intron 1 and 2 vs. intron 2) (**Fig. 4**). All but one ASO resulted in reduced *TRIBAL* abundance (although statistical significance was observed with only two ASOs in this suppression round) and, importantly, led to comparable suppression of several of the targets of interest. These findings confirmed that these ASOs specifically targeted *TRIBAL*.

### TRIB1 suppression partially phenocopies *TRIBAL* suppression in HepaRG and hepatocytes

TRIB1 is another important regulator of lipid metabolism. Its proximity to *TRIBAL* suggests that *TRIBAL* and TRIB1 may be functionally intertwined, as observed for several lncRNAs and their proximal protein coding gene [17]. Indeed, we previously demonstrated that *TRIB1* suppression resulted in increased *TRIBAL* expression in primary hepatocytes [5]. Moreover, *TRIBAL* suppression was generally associated with lower *TRIB1* expression, although statistical significance was only reached with ASO2 in one series of experiments (**Fig 2, 4**). Thus, *TRIBAL* may promote *TRIB1* expression, in turn facilitating TRIB1 function. First, the functional overlap between *TRIBAL* and TRIB1 was explored using publicly available data of antagomir-treated hepatocytes.

Targeting *TRIB1* or *TRIBAL* in hepatocytes resulted in a significant reduction of the TOIs, indicative of functional convergence (**Fig. 5A**). Consistency with HepaRG was then tested by qRT-PCR, using *TRIBAL* ASO2 and two distinct TRIB1 ASOs in HepaRG cells. Results were overall consistent with the hepatocytes findings, with *TRIB1* suppression resulting in similar (TRIB1 ASO2) or greater (TRIB1 ASO1) suppression than *TRIBAL* ASO2 in our transcript panel (**Fig. 5B**). A notable exception was *CYP7A1*, which was responsive to TRIB1 suppression only in HepaRG cells.

### Undifferentiated but confluent HepaRG are unaffected by *TRIBAL* targeting

Although differentiation of HepaRG post-confluence is important for the complete acquisition and maximal expression of hepatocyte-like traits, undifferentiated HepaRG cells express several hepatocyte-specific genes [6]. Consistent with a possible function, *TRIBAL* was expressed at levels comparable to differentiated cells, although considerable inter-experimental variation, not unique to HepaRG cells, was evident (**Fig S3**). To examine whether the *TRIBAL* function required complete differentiation, suppression was repeated in confluent but undifferentiated cells (10-14 days post-seeding). Unlike differentiated HepaRG, *TRIBAL* targeting consistently failed to reduce *TRIBAL* abundance (**Fig. 6A**). Indeed, *TRIBAL* expression was occasionally increased. By contrast, *TRIB1* silencing resulted in cognate suppression, confirming efficient cellular targeting (**Fig. 6B**). We reasoned that the considerable variation in *TRIBAL* and TOI abundance might reflect a potential causal relationship. Indeed, when the TOI values were plotted relative to *TRIBAL* abundance (normalized to the NT values), a clear pattern emerged: whereas most of the TOIs were not correlated to *TRIBAL* abundance. *TRIB1* levels uniquely clustered around 1, suggesting changes in *TRIBAL* and *TRIB1* were correlated (**Fig. 6C**). Regressing *TRIB1* against *TRIBAL* relative expression confirmed a monotonic relationship (**Fig. 6D**). Thus, although *TRIBAL* does not appear to be functional in undifferentiated HepaRG cells or may not be effectively targeted, these findings suggest that *TRIB1* and *TRIBAL* are co-regulated. Moreover, since *TRIB1* suppression did not affect *TRIBAL* abundance (Fig. 6B), the correlation is likely driven by *TRIBAL*, suggesting that *TRIBAL* can regulate *TRIB1* expression in that model system; however, this regulatory axis is insufficient to impact TOI expression.

### Comparison of *TRIBAL-*suppressed hepatocytes and HepaRG transcriptomes by Gene Set Enrichment Analysis

A transcriptomic investigation was then conducted to clarify the role of *TRIBAL* in primary hepatocytes and HepaRG cells and to delineate the functional footprints of TRIB1 and *TRIBAL*. To this end, *TRIBAL*- and *TRIB1*-ASO2-treated HepaRG samples were selected for transcriptome-wide expression analysis by microarrays. First, a comparison was made between *TRIBAL*-suppressed HepaRG cells and hepatocytes at the transcript level. *TRIBAL* suppression in HepaRG resulted in ∼ 5 times fewer nominally significant hits than in hepatocytes, suggesting either the possible loss of *TRIBAL* targets in HepaRG or lower quality data (**Fig. S4A**). Importantly, approximately half of the nominal HepaRG transcripts were also significant in Hepatocytes, indicating considerable convergence (p = 0.001, Jaccard similarity test).

To translate these differences into a biological understanding, the transcriptomes were then clustered to biological ontologies using Gene Set Enrichment Analysis (GSEA). The previously reported hepatocyte analysis was repeated to account for updates to the pipeline and dataset. The comparison of FDR significant terms revealed extensive overlap, predominantly in metabolic processes (**Fig. S4B, Tables S1-S2**).

Notably, the effects on the most impacted pathways were predominantly suppressive (negative effect size) in both models, indicating that these processes are curtailed in *TRIBAL*-targeted cells (**Table 1**). Examining terms specific to either cell type revealed growth and proliferation-related ontologies in HepaRG cells, whereas more metabolic ontologies were observed in primary hepatocytes (**Table S3**). Within shared ontologies, term clustering revealed a network of six terms centered on phospholipid metabolism (**Fig. S4C**).

**Table 1.** Comparison of shared and highly FDR significant ontologies in Hepatocytes and HepaRG treated with *TRIBAL* ASO2. Table showing shared and FDR highly significant (q<0.01) ontologies identified by GSEA in HepaRG and hepatocytes. Organized by incremental effect size in hepatocytes. NES, normalized effect size. For a complete list of impacted ontologies, see Tables S1 and S2.

|  | Hepatocytes |  | HepaRG |  |
| --- | --- | --- | --- | --- |
|  | NES | FDR | NES | FDR |
| **alcohol metabolic process** | -3.67 | 0 | -2.592 | 2.7E-03 |
| **fatty acid metabolic process** | -3.64 | 0 | -2.988 | 0 |
| **olefinic compound metabolic process** | -3.59 | 0 | -2.506 | 5.2E-03 |
| **steroid metabolic process** | -3.26 | 1.1E-04 | -2.388 | 6.4E-03 |
| **secondary metabolic process** | -3.14 | 9.4E-04 | -2.354 | 8.2E-03 |
| **sulfur compound metabolic process** | -3.09 | 1.4E-03 | -2.827 | 0 |
| **cellular ketone metabolic process** | -3.09 | 1.3E-03 | -2.355 | 9.0E-03 |
| **nucleoside bisphosphate metabolic process** | -2.91 | 7.8E-03 | -2.643 | 2.6E-03 |

### Transcriptome-wide impacts of *TRIB1* and *TRIBAL* suppressions in HepaRG

The functional interplay between *TRIBAL* and *TRIB1* was then examined by comparing their suppression in HepaRG cells. At the transcript level, nearly half of the *TRIBAL* ASO2-impacted transcripts were also nominally affected in *TRIB1* ASO2-treated cells (**Fig. S5A**). GSEA of *TRIB1*-targeted HepaRG cells identified 47 FDR-significant ontologies, of which 18 were similarly impacted in *TRIBAL*-suppressed cells (**Fig. S5B, Table 2, Table S4**). This shared group included several core metabolism terms, some of which showed a robust statistical association, including lipid and steroid-related ontologies (**Table 2, Table S5**). To complement these findings, GSEA was repeated using the *TRIB1* (rather than the corresponding CTLASO) expression values as the *TRIBAL* background, which the concurrent suppression design permitted. By aggregating transcript-level differences between *TRIB1-* and *TRIBAL-*targeted populations, we reasoned that this approach should enable the estimation of the relative contribution of *TRIB1* and *TRIBAL* to any given ontology. Clustering identified 11 FDR-significant enriched categories, including a single metabolic ontology, “glucose-6-phosphate metabolic process”, which was negatively enriched relative to *TRIB1*, suggesting that these functions are more inhibited in *TRIBAL-*suppressed cells (**Table S6**). In contrast, *TRIBAL* suppression was associated with the positive enrichment of a few FDR-significant terms linked to inflammation (“response to type II interferon,” “humoral immune response,” and “response to protozoan”), consistent with the establishment of a more inflammatory environment. Importantly, most of the major ontologies identified in the individual GSEA analyses were not significantly different in this analysis, indicating a generally concordant impact.

**Table 2.** Biological processes enriched in *TRIBAL*- and TRIB1-suppressed HepaRG cells. Impacted transcripts were mapped by GSEA to biological ontologies (non-redundant) via WebGestalt. FDR significant biological ontologies common to *TRIBAL*- and TRIB1-suppressed cells are shown below, organized by increasing effect size (HepaRG values). For a complete list of impacted ontologies in *TRIBAL* and TRIB1-suppressed cells, see Tables S2 and S3.

|  | HepaRG<br>*TRIBAL* |  | HepaRG<br>TRIB1 |  |
| --- | --- | --- | --- | --- |
|  | NES | FDR | NES | FDR |
| **fatty acid metabolic process** | -3.0 | 0 | -3.4 | 0 |
| **sulfur compound metabolic process** | -2.8 | 0 | -2.3 | 1.2E-02 |
| **nucleoside bisphosphate metabolic process** | -2.6 | 2.6E-03 | -2.9 | 0 |
| **alcohol metabolic process** | -2.6 | 2.7E-03 | -2.8 | 0 |
| **olefinic compound metabolic process** | -2.5 | 5.2E-03 | -2.3 | 1.3E-02 |
| **steroid metabolic process** | -2.4 | 6.5E-03 | -2.8 | 0 |
| **cellular ketone metabolic process** | -2.4 | 9.0E-03 | -2.3 | 1.2E-02 |
| **fatty acid derivative metabolic process** | -2.3 | 1.1E-02 | -2.6 | 1.7E-03 |
| **small molecule catabolic process** | -2.3 | 1.2E-02 | -2.2 | 1.9E-02 |
| **hormone metabolic process** | -2.2 | 1.3E-02 | -2.0 | 3.2E-02 |
| **isoprenoid metabolic process** | -2.2 | 1.6E-02 | -2.2 | 2.1E-02 |
| **organic acid biosynthetic process** | -2.2 | 1.7E-02 | -2.4 | 1.2E-02 |
| **platelet-derived growth factor receptor signaling pathway** | -2.2 | 1.8E-02 | -2.0 | 3.2E-02 |
| **post-embryonic development** | -2.1 | 2.3E-02 | -2.2 | 1.9E-02 |
| **lipoprotein metabolic process** | -2.0 | 3.6E-02 | -2.2 | 1.8E-02 |
| **smoothened signaling pathway** | -2.0 | 3.6E-02 | -1.9 | 4.4E-02 |
| **lipid modification** | -2.0 | 3.4E-02 | -2.2 | 1.8E-02 |
| organic hydroxy compound<br>catabolic process | -2.0 | 3.8E-02 | -1.9 | 4.9E-02 |

### Transduced *TRIBAL*1 cannot rescue *TRIBAL* suppression

We previously demonstrated that *TRIBAL1* overexpression had no impact on primary hepatocytes, implying that transduced *TRIBAL1* might be dysfunctional or insufficient [1]. As with primary hepatocytes, expression of the gene panel was not affected in HepaRG transduced with *TRIBAL1* (**Fig S6**). To test whether *TRIBAL1* is functional, a HepaRG pool stably expressing *TRIBAL1* was targeted using the intron-specific ASO2, thus not targeting transduced *TRIBAL1*. We reasoned that the presence of *TRIBAL1* should offer protection from ASO2 only if *TRIBAL1* is functional. Indeed, whereas the endogenous *TRIBAL* in the transduced PLVX controls was highly sensitive to ASO2, resulting in greater than 90% suppression of *TRIBAL*, overexpressed *TRIBAL*1 was unaffected (**Fig S7**). Importantly, *TRIBAL1* did not protect against ASO2. As off-target effects of ASO2 were deemed improbable, given the consistent impact of several other cognate ASOs (**Fig 4**), these experiments suggested that overexpressed *TRIBAL1* was dysfunctional.

### Endogenous, but not transduced *TRIBAL*, is resistant to RNAseR

We hypothesized that *TRIBAL1* dysfunction may stem from improper folding or maturation. The native expression locus may impart unique features that the lentiviral insertion sites may not provide. Structural differences were probed using RNAseR, a structure-sensitive RNA exonuclease. Strikingly, recombinant *TRIBAL1* was highly sensitive to RNAseR, on par with the PPIA mRNA. By contrast, native *TRIBAL*, like U1, a highly structured snRNA, showed a near-complete resistance to RNAseR (**Fig 7**). Thus, the native locus may be key in conferring functionality to *TRIBAL*, possibly by imparting unique folding attributes.

## Discussion

*TRIB1* and *TRIBAL* are both needed to support hepatocyte function. Like *TRIBAL*, we previously demonstrated that *TRIB1* suppression also reduced *HNF4A* expression and function in primary hepatocytes [18]. Given the central role of HNF4A in establishing and maintaining liver function, substantial functional epistasis between *TRIB1* and *TRIBAL* is to be expected. Moreover, the subset of transcripts populating our TOI list, defined in our previous work based on the effects of *TRIBAL* suppression, was also inhibited by *TRIB1* suppression. Our transcriptomic enrichment analyses are also consistent with this convergence, although unique contributions were identified, suggesting incomplete epistasis. However, one must interpret these “unique” contributions with caution, as the residual expression may suffice to ensure partial functionality. Thus, these clustering approaches likely underestimate the true participation of *TRIBAL* and TRIB1.

Future investigations will examine the possibility of mechanistic interplays between *TRIBAL* and TRIB1. LncRNAs may operate locally and act as enhancer components in disease [19,20]. In line with this model, the *TRIBAL* locus has been hypothesized to regulate liver function through TRIB1, whose role in hepatic function is well established. However, recent assessments of the relative contribution of proximal protein-coding and non-coding genes using transcriptome-wide interventions necessitate a reassessment of this model. Focusing on essentiality criteria, 778 lncRNAs that were essential for proliferation in at least 1 of 5 cell lines were identified [21]. Remarkably, these were prone to operate independently of neighboring protein-coding genes, highlighting their functional independence.

Here we used differentiated 2D HepaRG cultures as models for *TRIBAL* studies. Using an algorithm prioritizing a panel of liver-specific transcripts, Kim et al. have shown 3D and 2D cultures of HepaRG cells to have 59% and 41% liver similarity, demonstrating that both models are good but imperfect approximations of hepatocytes; however, both were superior to hepatocyte-like cells derived from iPSC in that regard [15]. Interestingly, HepaRG cells grown in 2D can differentiate (and transdifferentiate) into hepatocyte- and biliary-like cells in approximately equal proportions [22]. It will be interesting to examine the implication of *TRIBAL* in this process since lncRNAs can regulate pluripotency and differentiation[2]. The role of *TRIBAL* in biliary cells remains unexplored, and its absence may contribute to the weaker impact of *TRIBAL* ASOs on HepaRG cells compared to primary hepatocytes, as identified by our enrichment analyses. Alternatively (or in addition to), the more modest effect of *TRIBAL* suppression in HepaRG cells could be due to the incomplete acquisition of hepatocyte traits.

It is unclear how and why undifferentiated HepaRG cells resist *TRIBAL* suppression. Control *TRIB1* suppression was effective and reproducible, arguing that the ASOs were delivered appropriately and that the RNA degradation machinery was functional. Moreover, following *TRIBAL* suppression, TRIB1 exhibited a significant correlation with *TRIBAL* expression, suggesting that *TRIBAL* abundance was accurately measured. *TRIBAL* targeting was associated with highly variable transcript abundance rather than having no impact. This apparent noise may reflect genuine biological variation, possibly elicited by *TRIBAL* targeting. It is also unclear why *TRIBAL* is not reproducibly suppressed in these cells.

Although HepG2 and HuH-7 cells were inadequate models for the study of *TRIBAL*, the locus affected cell proliferation in these cells, as revealed by the selection of unedited or minimally edited populations. Since *TRIBAL* suppression did not reduce HepG2 proliferation (HuH-7s were not tested), we speculate that the locus may play additional roles independent of its transcript. This may include supporting TRIB1 expression, which promotes cell growth in both cell models, through enhancer-like effects [23,24]. The transformation of HepG2 and HuH-7 may also activate pathways that neutralize or antagonize *TRIBAL* metabolic functions.

Interestingly, native and recombinant *TRIBAL* transcripts differed fundamentally in their sensitivity to RNAseR. This may underlie the inability of the transduced *TRIBAL1* to protect from *TRIBAL* suppression or single-handedly regulate TOI expression. The proximal reasons for this difference are unknown. They are likely unrelated to transcript sequence, as *TRIBAL1* corresponds to the predominant form in HepG2 cells and is the only form identified in primary hepatocytes. Rather, we propose that its synthesis from the native locus is critical for accurate folding and impart function. *TRIBAL* emerging from its native locus may mature differently from the transduced transcript through the participation of locus-enriched chaperones and/or epi-transcriptome chemical modifications. Our understanding of RNA methylation is still in its infancy, but evidence suggests it helps define lncRNA function [25].

## Acknowledgment

This work was supported by the Canadian Institutes of Health Research (CIHR) FDN-154308 (RM).

---

**Figure S1.**
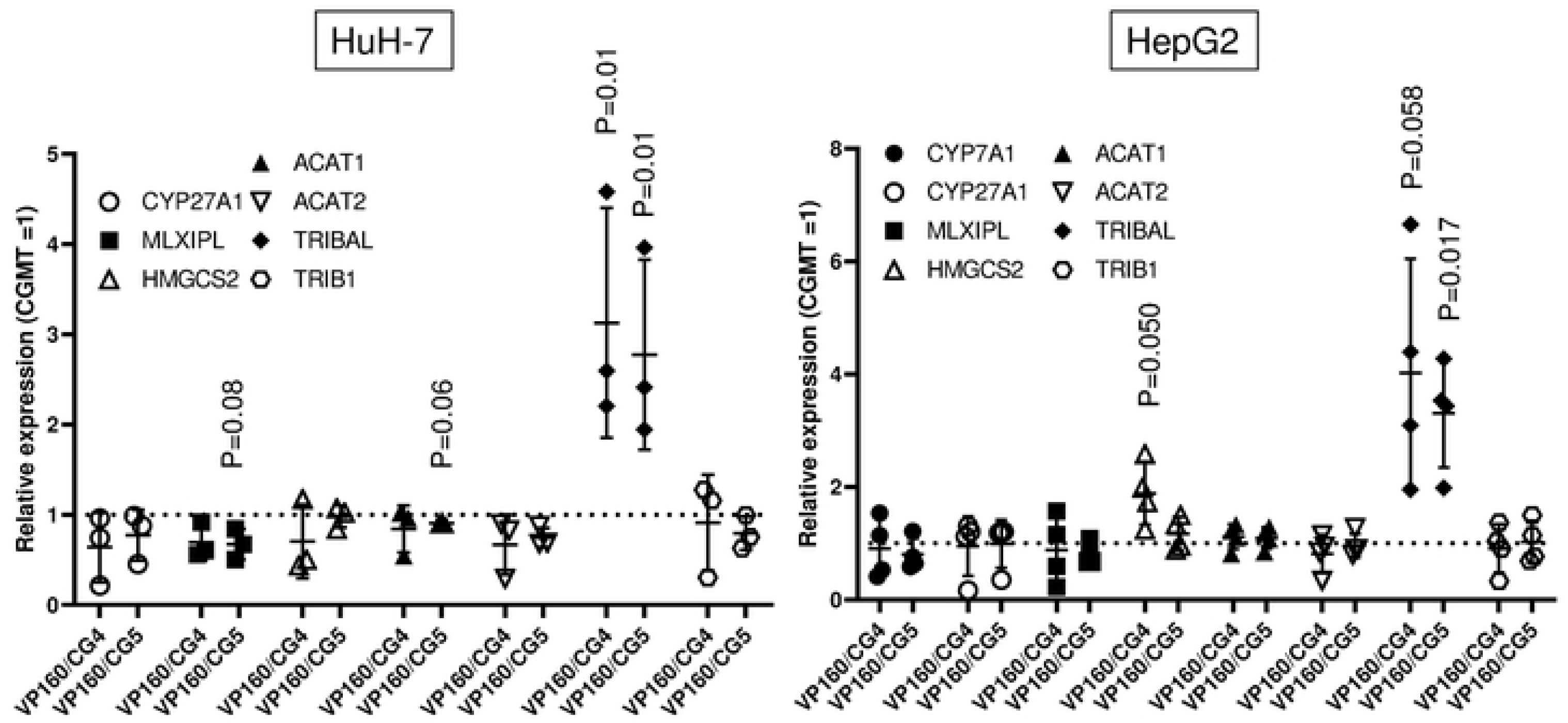
TRIBAL activation by dCRISPRa in HuH-7 and HepG2 does not impact the transcript panel significantly. Cells were transfected *tor* 72 h with a plasmid encoding dCAS9-VP160 and sgRNA sequences targeting the promoter region of TRIBAL. Each point represents a biological replicate. Statistical significance was assessed using a one-sample t-test using a theoretical control value of 1 in Prism. Nominal P values approaching (p≤to 0.1) or surpassing nominal significance are shown. CYP7A1 abundance in HuH-7 cells was too low to be reproducibly measured.

**Figure S2.**
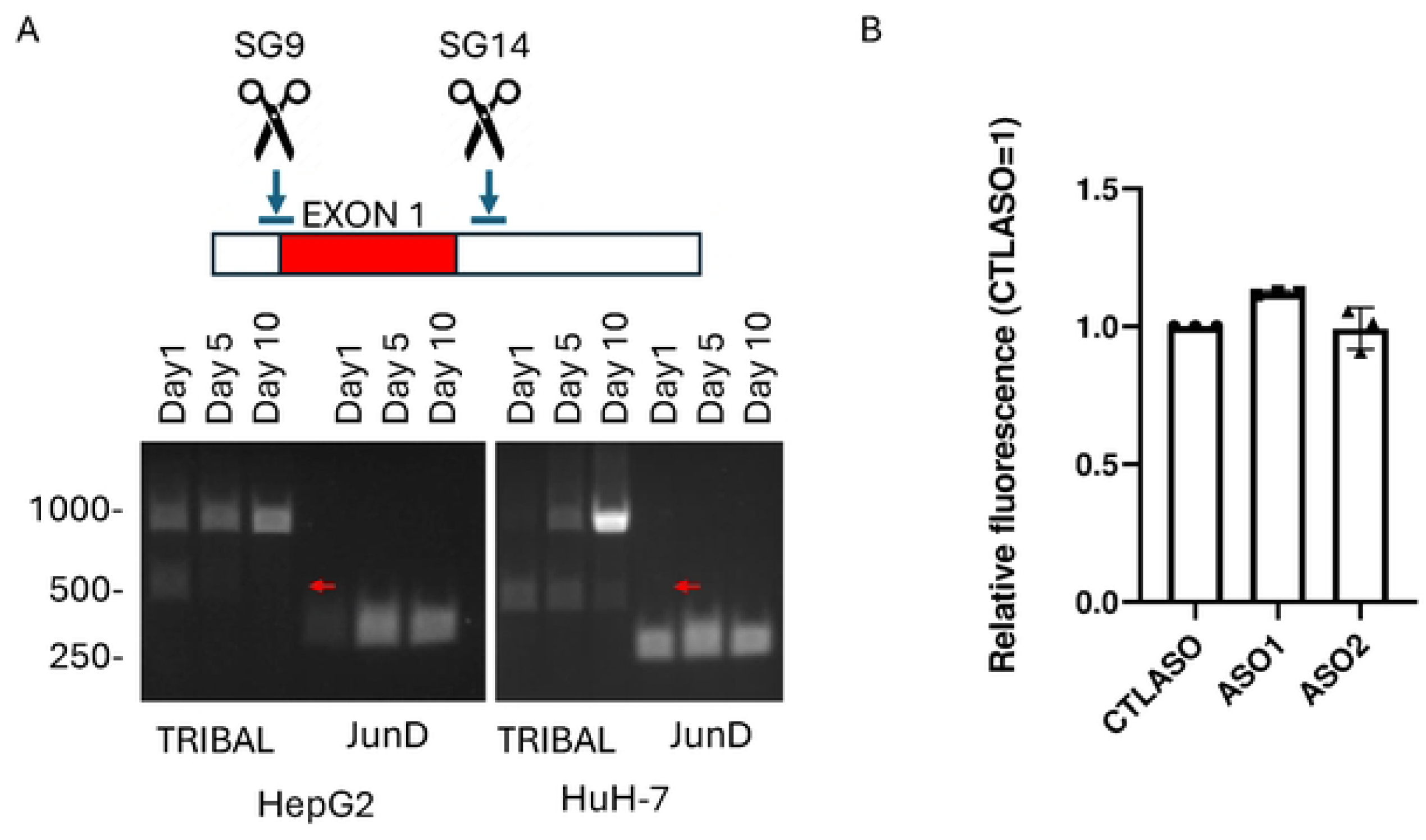
Loss of TRIBAL is deleterious and selected against in HepG2 and HuH-7. A, TRIBAL was targeted with CRISPR and 2 sgRNAs flanking exon 1 (sg9 and sgl 4). The schema of cg positions is shown at the top. The red arrow points to the position of the deleted allele. The experiment was repeated thrice (HepG2) or twice (HuH-7), with similar results. DNA from whole cell lysates was amplified with primers flanking the TRIBAL sgRNA cognate sites or targeting Juno (positive control). B, Impact of TRIBAL ASOs on HepG2 cell proliferation. A one-tailed I-test, performed under the hypothesis that TRIBAL suppression should reduce cell proliferation, showed no significant difference.

**Figure S3.**
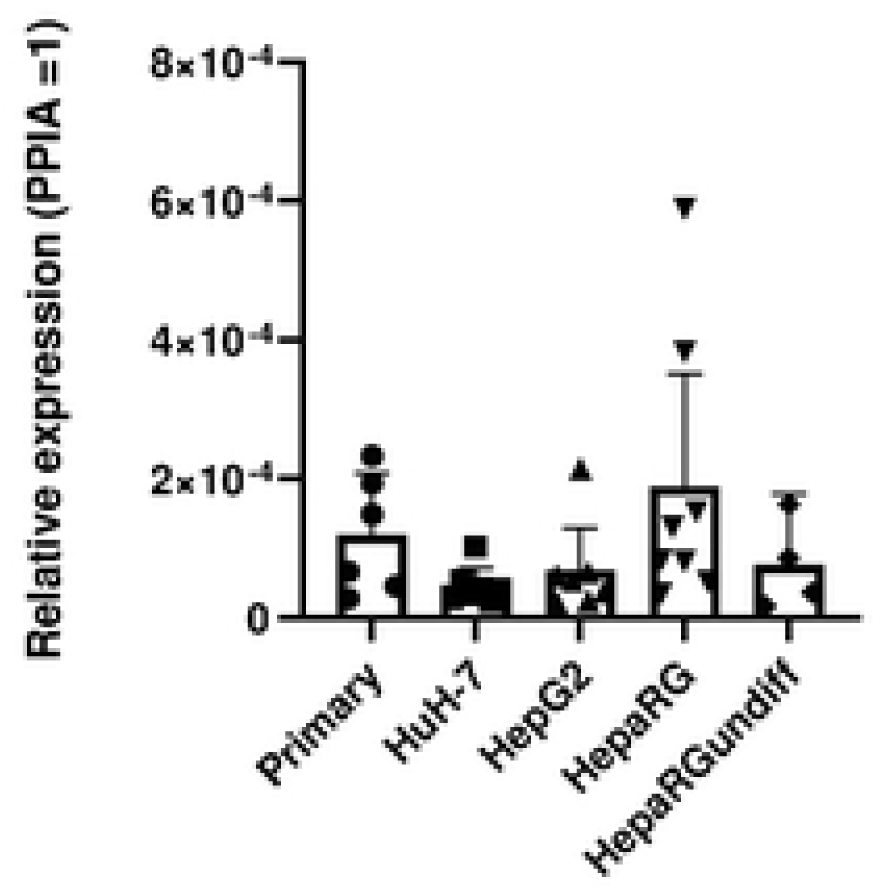
TRIBAL expression in hepatocyte cell models. TRIBAL expression was measured in cell models treated with the CTLASO tor 72 h. Each point represents a distinct biological replicate. Expression is expressed relative to PPIA. To simplify visualization, error bars below the mean are not shown.

**Figure S4.**
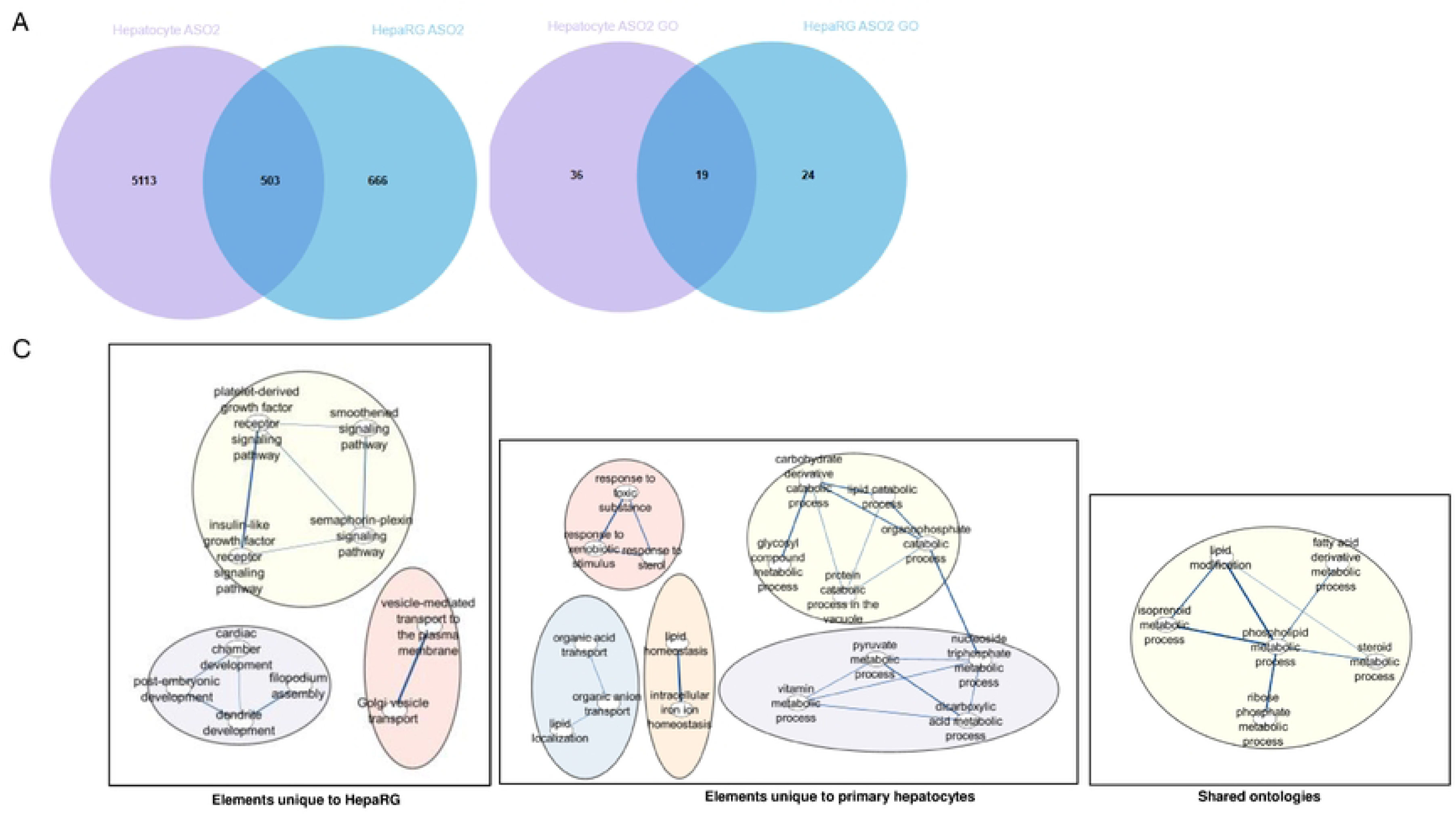
Functional consistency between HepaRG and hepatocytes treated with TRIBAL AS02. A, Venn diagram of nominally impacted Transcript IDs (mapped 10 Entrez Genes) in HepaRG cells and hepatocytes. B, Venn diagram of FDR significant Gene Ontology terms identified by GSEA in TRIBAL-suppressed in HepaRG cells and hepatocytes. HepaRG (magenta) and hepatocytes (blue). C, the ontologies from B were clustered with Revigo and exported into Cytoscape for visualization. Singletons were removed to aid visualization. A complete list is shown in Tables S-1S2.

**Figure S5.**
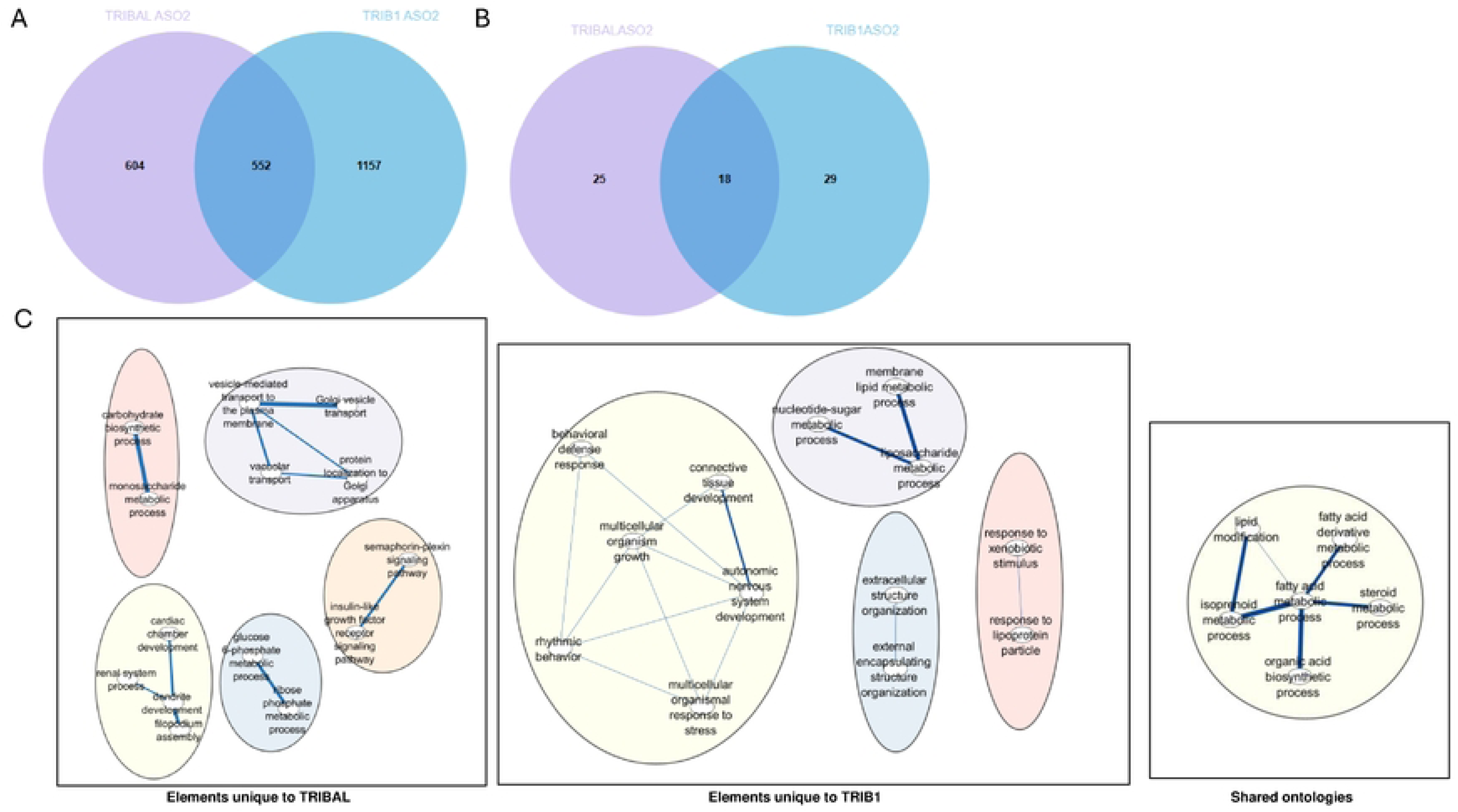
Functional overlap between TRIBAL and TRIB1 in HepaRG cells. A, Venn diagram of nominally impacted Transcript IDs (mapped to Entrez Genes) in HepaRG cells targeted with TRIBAL or TRIB1AS02. B, Venn diagram of FDR-significant Gene Ontology term sidentified by GSEA in HepaRG cells targeted with TRIBAL or TRIB1AS02. TRIBAL (magenta) and TRIB1 (blue). C, categories from B were clustered with Revigo and exported into Cytoscape for visualization. Singletons were removed to aid visualization. See Tables S2-S3 *tor* the complete list of ontologies

**Figure S6.**
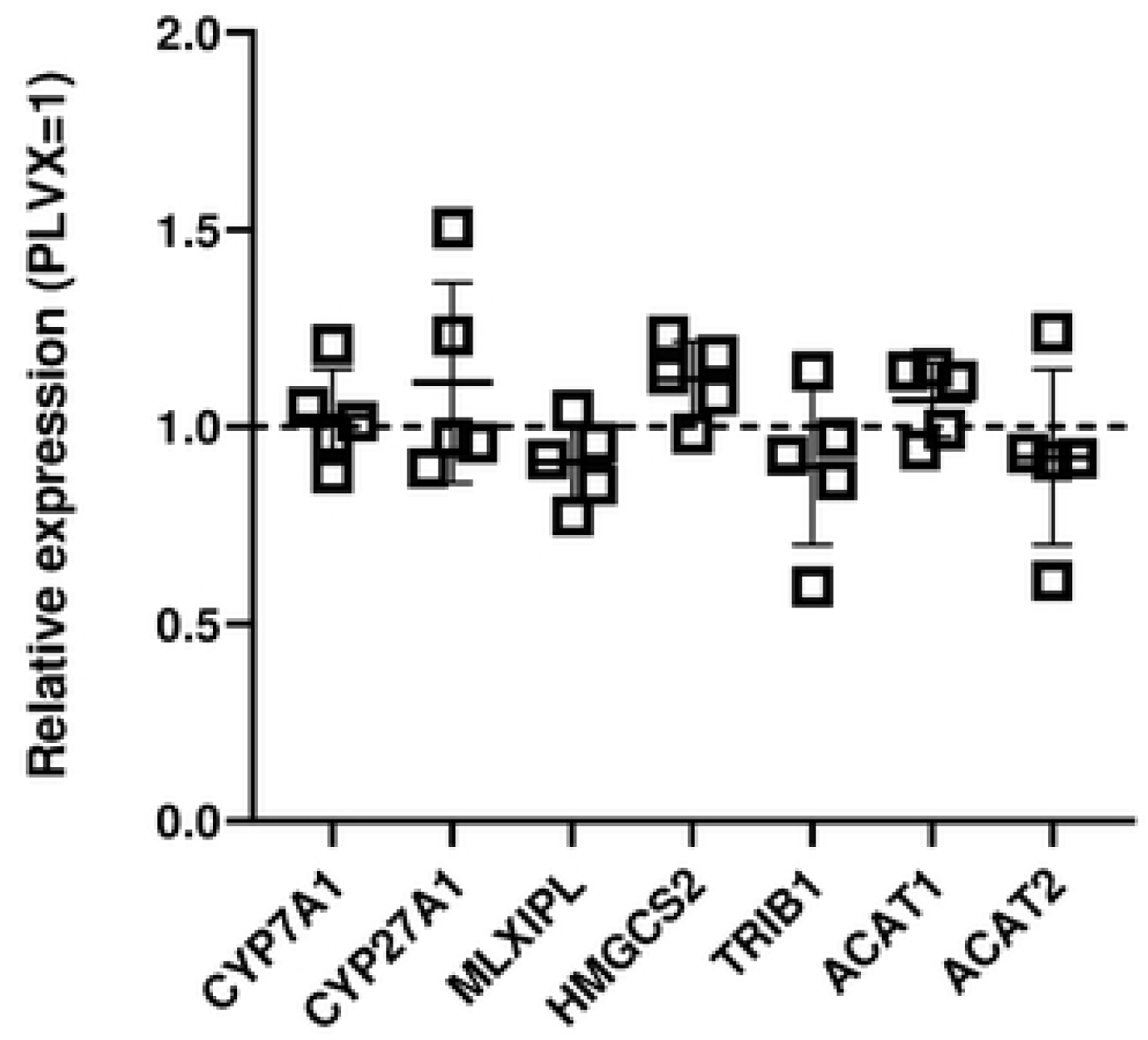
Impact of TRIBAL1 Transduction on HepaAG Transcripts of Interest. HepaRG cells were transduced a PLVXTRIBAL1 or PLVX for 72 h. RNA was then isolated and quantified by qRT-PCR. TRIBAL was upregulated by 6700 ± 4600(S.D.). Values were internally normalized to PPIA and are expressed relative to the values from the PLVX transduced controls. Differences were not statistically different from the PLVX values (one sample I-test using a theoretical control value of 1), except for TRIBAL (p•0.032).

**Figure S7.**
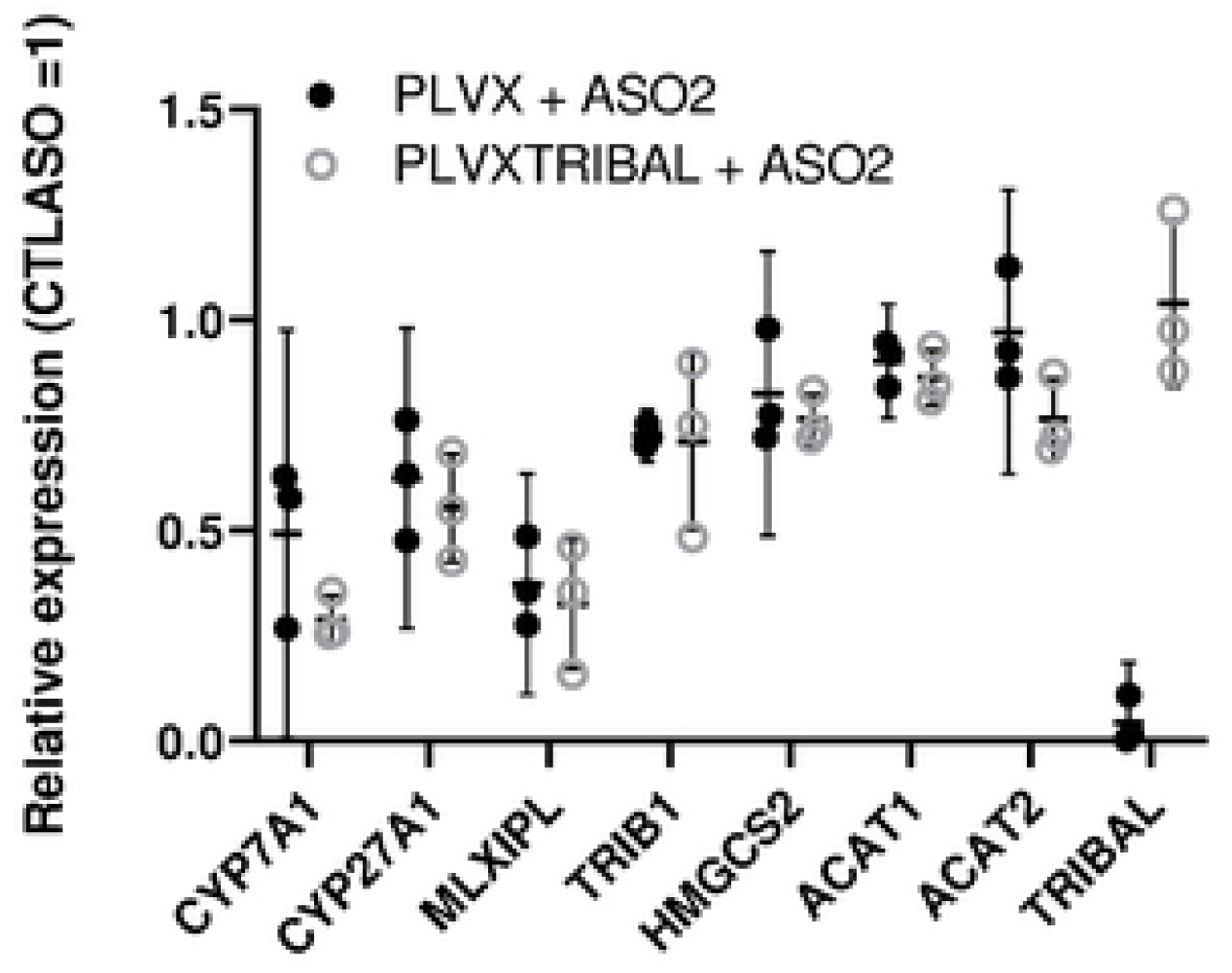
TRIBAL OE in HepaRG cannot protect from the Impacts of TRIBAL suppression. HepaRG cells were transduced with PLVXTRIBAL1 or PLVX for 48 h before treatment with AS02 or CTLASO for 72 h. RNA was then isolated, converted to cDNA and analyzed for the indicated targets by qRT-PCR. TRIBAL transduction resulted in a 460-fold (± 350,S.D.) increase in TRIBAL abundance. Each point is a biological replicate.

